# Cross-Continental Analysis of Vampire Bat Betaherpesvirus Reveals Limited Interference Among Strains and Local Geographic Spread

**DOI:** 10.64898/2026.05.26.727811

**Authors:** Haris Malik, Richard Orton, Anushka Ramjag, Alice Broos, Aline Campos, Andre Witt, Anne Lavergne, Carlos Tello, Christine Carrington, Daniel J. Becker, Elsa Cárdenas-Canales, Erich Fischer, Gerald G. Carter, Luke Rostant, Megan Griffiths, Molly C. Simonis, Nancy B. Simmons, Nicholas Mohammed, Rosana Huff, Sebastian Stockmaier, Michael Jarvis, Laura Bergner, Daniel G. Streicker

## Abstract

*Desmodus rotundus* Betaherpesvirus (DrBHV) is a candidate vector for a transmissible vaccine targeting the circulation of rabies virus within its vampire bat reservoir. Studies assessing the potential for DrBHV as a vector have not considered its geographic range, potential for transboundary spread or how the diversity of wildtype DrBHV in natural bat populations might impede the spread of a modified vaccine strain. Here, by sequencing DrBHVs from vampire bats spanning 15 regions across 7 rabies-affected countries in Latin America and the Caribbean, we characterise the continent-scale distribution of DrBHV diversity and demonstrate widespread and apparently unconstrained co-infection. DrBHV occurred in all regions, forming a monophyletic clade consistent with a single introduction to vampire bats or host–virus co-speciation rather than frequent host switching. Phylogeographic analyses revealed cross-boundary spread that was predicted by geographic proximity. We identified 50 putative DrBHV strains and 79% of DrBHV-infected bats harboured multiple strains. No strains were over- or under-represented in co-infection and co-infections occurred proportionately to local strain prevalence. Whether strains circulated in given populations was predominantly driven by the geographic proximity of other populations containing that strain. The evolutionary relatedness of strains constrained neither rates of co-infection within individuals nor whether strains co-circulated within regions, suggesting vaccine vectors might be locally sourced rather than requiring imported, divergent strains to ameliorate interference. These results support the viability of DrBHV-vectored vaccines for mitigating rabies virus across Latin America and the Caribbean, despite widespread circulation of wildtype viruses.

## Introduction

Rabies virus causes an acute, almost universally lethal zoonosis in all mammals. In most Latin American and some Caribbean countries, common vampire bats (*Desmodus rotundus*) are the primary animal source of human rabies mortality, outnumbering cases arising from dogs and other vertebrate reservoirs ^1^. Vampire bat-transmitted rabies virus (VBRV) further causes significant losses in the livestock industry, exceeding ∼US$30 million per year in cattle mortality alone ^2^, and disproportionately affects subsistence farmers, contributing to perpetual poverty ^3^. Current interventions have proved largely ineffective ^3^. Livestock vaccination is typically carried out reactively to outbreaks and does not disrupt transmission in wild bat populations. Culling of bats has been used since the 1970s. While culls generally reduce the incidence of bat bites, benefits for rabies prevention remain unclear ^4–8^. In Peru, culling was associated with higher rabies seroprevalence within *D. rotundus* populations ^9^, failed to reduce the incidence of rabies in livestock and accelerated the spatial spread of rabies when implemented reactively to outbreaks ^10^. Moreover, culling as currently implemented has been insufficient to halt ongoing viral invasions into historically rabies-free areas ^11–14^. These shortcomings underline the requirement for novel interventions.

Mass vaccination of animal reservoirs has been an effective control (and elimination) strategy for carnivore-mediated rabies ^15–18^. By reducing viral circulation within reservoir hosts, mass vaccination prevents spillover at the source. However, neither parenteral nor bait-based mass vaccination strategies which have been adopted for carnivore vaccination are suitable for bats because of their large population sizes, isolated roosts, elusive behaviour and specialised diets ^3^. An emerging theoretical solution to this scalability challenge is transmissible vaccines (TVs). Modern TV approaches propose to utilise naturally benign, endemic viruses as recombinant viral vectors (RVVs) that would express immunogenic antigens of a target virus without attenuating their ability to spread autonomously through, and therefore immunise, wildlife populations ^19–21^. Betaherpesviruses are strong candidate TV vectors, owing to their generally benign infection and capacity to express foreign inserts ^20–27^.

The best-developed example in bats is *D. rotundus* Betaherpesvirus (DrBHV), first detected via metagenomic screening of pooled oropharyngeal swabs from *D. rotundus* populations in Peru ^28^. Subsequent work demonstrated that bats experience lifelong infections characterised by latency and periodic reactivation, a trait that could extend the window for onward transmission of a vaccine-derived strain or provide a self-boosting mechanism ^29^. Longitudinally monitored bats also acquired additional strains throughout their lifetimes (‘superinfection’), suggesting that a vaccine strain might also infect hosts with concurrent or past wildtype (WT) infections ^30^. Importantly, WT infection in vampire bats is not associated with evident pathology, supporting its suitability as a non-attenuated vector. Finally, epidemiological models based on patterns of infection in wild bats suggested that a DrBHV-based TV could achieve >80% coverage in *D. rotundus* populations after a single inoculation, reducing the magnitude, frequency and duration of VBRV outbreaks by 50–95% ^29^.

Existing data supporting DrBHV as a TV vector originate from Peru, representing a relatively narrow part of the natural range of *D. rotundus*, which spans most of Central and South America and parts of North America ^8,31–34^. As such, the continent-wide diversity of DrBHV lineages, how the virus may spread at large geographic scales (e.g., between countries) or how local ecological conditions influence infection prevalence remain unknown. These uncertainties underpin both safety and regulatory planning for eventual TV deployments and the technical feasibility and performance of future vaccines. For example, the potential for TVs to spread between administrative boundaries creates shared risks and benefits which may require harmonised surveillance, data sharing and predefined escalation criteria across jurisdictions. The expected extent and determinants of transboundary movements could inform the geographic extent of coordination required. From the perspective of technical feasibility and vaccine performance, reduced DrBHV prevalence in parts of the *D. rotundus* range could imply low potential for vaccine spread and, therefore, the need for higher deployment efforts. Perhaps most importantly, competition with WT strains is a foundational concern with using endemic viruses as TVs ^35^. Earlier work showed that vampire bats harbour multiple DrBHV strains simultaneously; however, the ecological and evolutionary processes that shape co-infection within individuals and co-circulation at the population level could not be tested with the relatively low number of strains detected ^30^. In particular, it remains unclear how the genetic similarity of strains structures coexistence. For example, if strains are less likely to occur in regions where closely-related lineages already circulate, this might suggest competitive exclusion via cross-immunity. Similarly, among co-circulating strains, lower than expected frequencies of co-infection among closely-related strains within hosts would signal within-host competition or interference. If closely-related lineages are systematically less able to co-circulate or co-infect, this would suggest that resident WT strains could impede establishment and onward spread of a genetically similar vaccine backbone, making vaccine performance contingent on local strain composition or on the discovery of strains with exceptional superinfection capacity. Alternatively, establishing that very closely related strains can co-infect and co-circulate would support using locally-circulating viruses as vectors, which would eliminate concerns associated with introducing divergent strains from other regions to avoid inter-strain competition.

Against this backdrop, by analysing 229 *D. rotundus* samples collected across Latin America and the Caribbean, we aimed to: (1) determine DrBHV prevalence and phylogenetic structure across the range of *D. rotundus*; (2) identify inter-regional transmission routes for DrBHV and the spatial or ecological predictors of these transitions; (3) determine if genetic similarity shapes the regional co-circulation of DrBHV strains; (4) test for patterns of genetic similarity between co-infecting strains; and (5) test for over- or under-representation of particular strains in multi-strain infected individuals. Collectively, this work demonstrates that transboundary spread of a DrBHV vectored vaccine would occur over small scales and suggests weak barriers to superinfection that might otherwise frustrate the use of DrBHV in rabies prevention.

## Results

### DrBHV is Common and Monophyletic across the Geographic Range of Vampire Bats

We used long-range PCR to measure DrBHV prevalence across 15 sampling regions from 7 countries, targeting a 3,919 bp genomic region that spanned 849 bp of the DNA polymerase gene (DPOL; UL54) at the 5′ end, the full ∼2991 bp glycoprotein B gene (gB; UL55), and 79 bp of the UL56 gene at the 3′ end. We carried out analyses at the regional level, with regions defined as either countries or first-level administrative subdivisions (depending on whether multiple first-level subdivisions were sampled within a given country) to maximise spatial resolution whilst maintaining sufficient sample sizes. Overall, 60.7% (139/229) of sampled vampire bats were infected with DrBHV; however, prevalence ranged from 26.3% to 100% across regions (Table 1), and from 26.3% to 84.6% across countries (Table 1, Fig. 1(B)). Substantial heterogeneity was evident within Peru and México – highlighting spatial variation in prevalence at finer geographic scales. It should be noted that some regions have low sample sizes (e.g. Loreto, Peru [n=6] and Amazonas, Peru [n=4]), so these prevalence estimates should be interpreted with caution.

**Figure 1:**
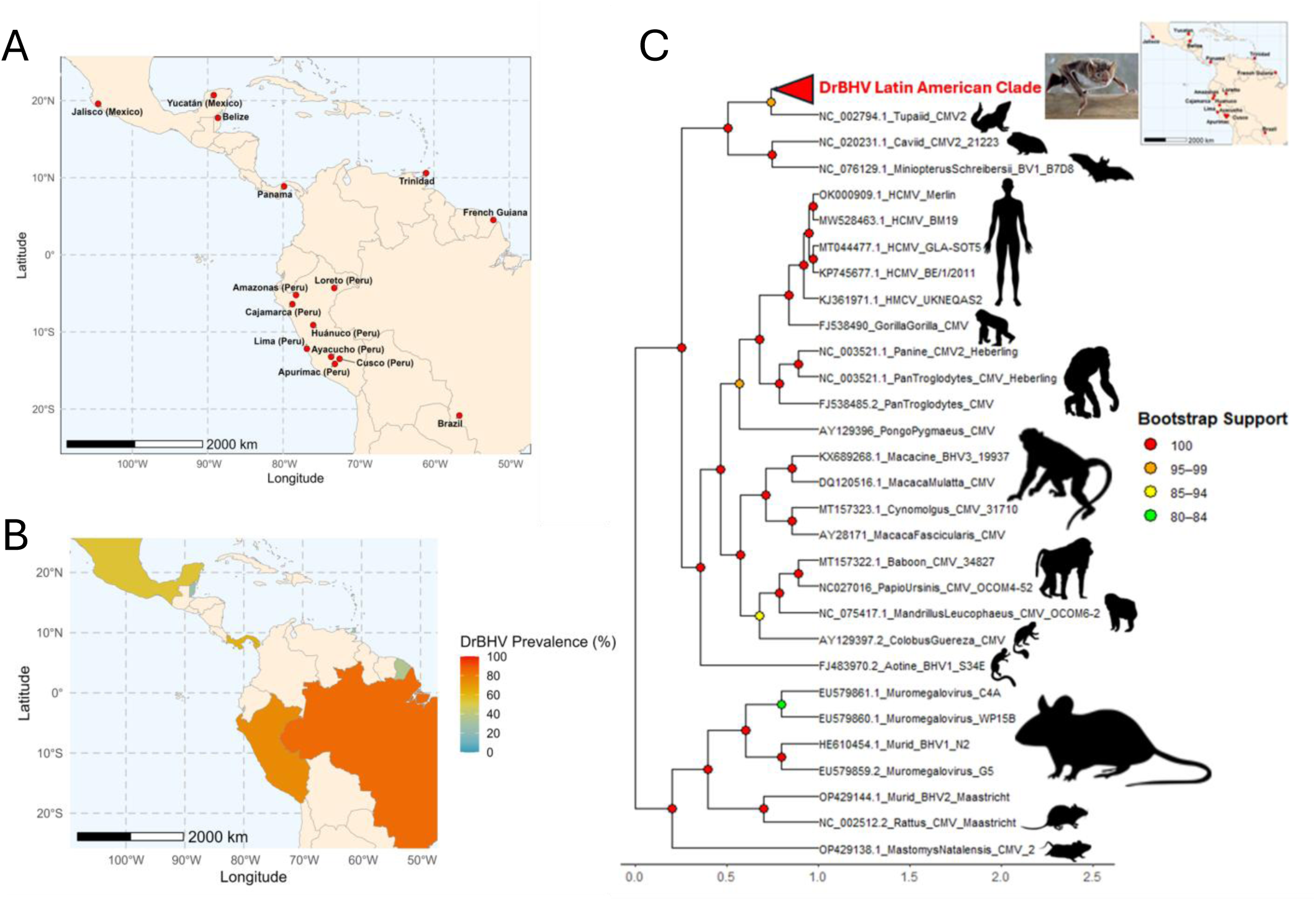
DrBHV is common and monophyletic across the geographic range of *D. rotundus*. **(A)** Shows sampling regions as red circles, covering México (Jalisco, Yucatán), Belize, Panamá, Trinidad, French Guiana, Brazil and Peru (Amazonas, Loreto, Cajamarca, Lima, Huánuco, Ayacucho, Apurímac, and Cusco). **(B)** Shows prevalence across countries using a heatmap layer projected onto a map of Latin America, calculated as the proportion of DrBHV-positive bats detected by PCR of a ∼3,919 bp region spanning part of the DPOL gene, the full gB gene, and part of the UL56 gene. Unsampled countries are coloured the same as the basemap layer. **(C)** Shows the maximum likelihood phylogeny of the same genomic region. This analysis includes all 627 *D. rotundus* sequences sampled across Latin America, in context with other homologous mammalian Betaherpesvirus sequences from GenBank (Table S3). For visualisation, the monophyletic clade containing all 627 DrBHV sequences was collapsed. The phylogeny was reconstructed by IQ-TREE2 using a GTR+F+R10 nucleotide substitution model, as selected by ModelFinder (lowest BIC), with 10,000 bootstrap replicates. The tree was rooted using KC465951.1_HHV-6A_GS (Human Herpesvirus 6A – Roseolovirus) as an outgroup, which was dropped for visualisation purposes. Coloured circular node points indicate bootstrap support categories. Animal silhouettes were added to the final tree to aid visualisation, obtained from PhyloPic.

**Table 1:**
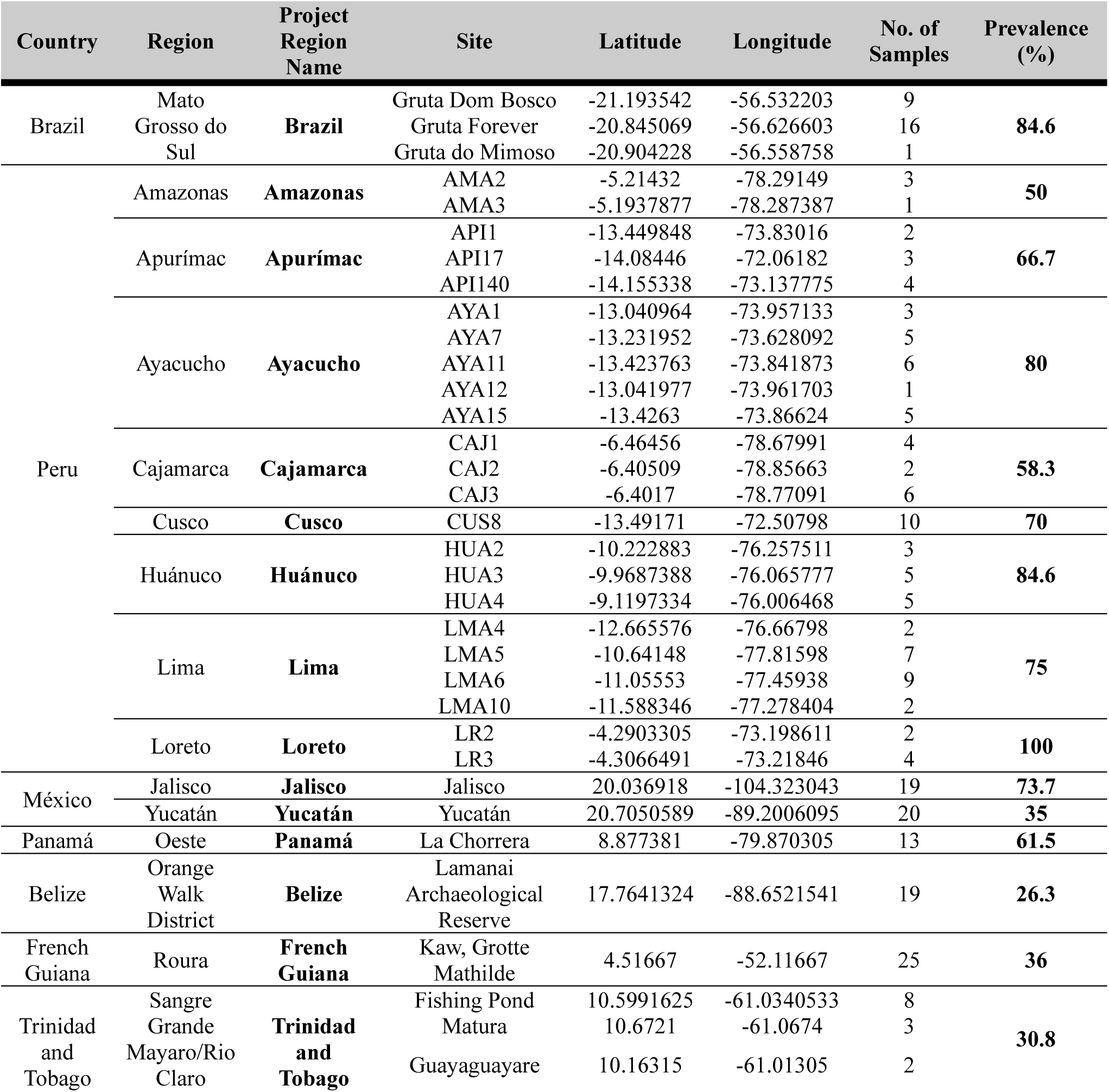
DrBHV prevalence across regions and spatial organisation of sampling regions. Prevalence of DrBHV in *Desmodus rotundus* across different regions in Latin America, based on successful amplification of a ∼3,919 bp region of the DrBHV genome flanking the entire glycoprotein B (gB) gene, as well as a portion of the viral DPOL and UL56 genes. Prevalence was calculated as the percentage of individuals testing positive in each region. The spatial organisation and geographical coordinates of sampling sites is also shown: Country → Region → Site. Sample sizes per site are also shown. Analyses were carried out at the regional level to maximise spatial resolution whilst maintaining acceptable sample sizes. The column *Project Region Name* indicates the names assigned to each region, or groups of regions, for downstream analyses.

To identify regional or biological variation in the prevalence of DrBHV, we fitted generalised linear mixed models (GLMMs) with DrBHV infection as a binary response variable, including administrative boundaries and host-related traits of bats as fixed effects, and capture year as a random effect. Due to metadata omissions, we excluded samples from French Guiana and Trinidad from this analysis. The most parsimonious GLMM after model selection retained only bat reproductive status, indicating that reproductively active individuals had 62% lower odds of detectable DrBHV infection compared to reproductively inactive individuals (β = −0.964, SE = 0.369, z = −2.62, p = 0.0089; Table S1). An additional GLMM including the interaction between sex and reproductive status was less parsimonious and the interaction did not improve model fit (ΔAICc = ∼3.45; likelihood ratio test [LRT]: χ²(2) = 0.55, p = 0.76; Table S2), indicating no evidence that the effects of reproductive status differed between male and female bats. Despite variation in raw prevalence among regions, region was not retained in any top-ranked model (Table S1), providing no support for significant regional variation in DrBHV prevalence.

Nanopore sequencing of DrBHV-positive amplicons from 139 infected bats generated 627 unique DPOL-gB sequences which formed a highly-supported monophyletic group (UFboot Support = 100) within the larger mammalian BHV tree (Fig. 1(C)). The maximum genetic divergence within the DrBHV clade was 9.24% (between Brazil19-C0 and Lima5-C2), whereas the smallest genetic distance between a DrBHV sequence (Jalisco8-C30) and the closest non-DrBHV sequence (NC_002794.1_Tupaiid_CMV2) was 36.32%. Together, these results indicate DrBHV as a single virus species found throughout the range of vampire bats that is highly divergent from other reported mammalian BHVs with publicly available DPOL-gB sequences.

### Regional Spread of DrBHV is Predicted by Geographic Proximity

To reconstruct inter-regional spread of DrBHV from the DPOL–gB sequence data, we developed a Bayesian phylogeographic model incorporating Bayesian stochastic search variable selection (BSSVS) to estimate support for transitions among regions. The resulting maximum clade credibility (MCC) phylogenetic tree revealed multiple well-supported DrBHV lineages circulating across Latin America and the Caribbean. Although several lineages contained sequences from multiple regions, sequences tended to cluster geographically within lineages, indicating substantial regional structure. The phylogeny revealed both region-specific clades, consistent with locally restricted or recently evolved endemic strains, and mixed-region clades, consistent with broader spatial spread of DrBHV lineages across the vampire bat range (Fig. 2A).

**Figure 2:**
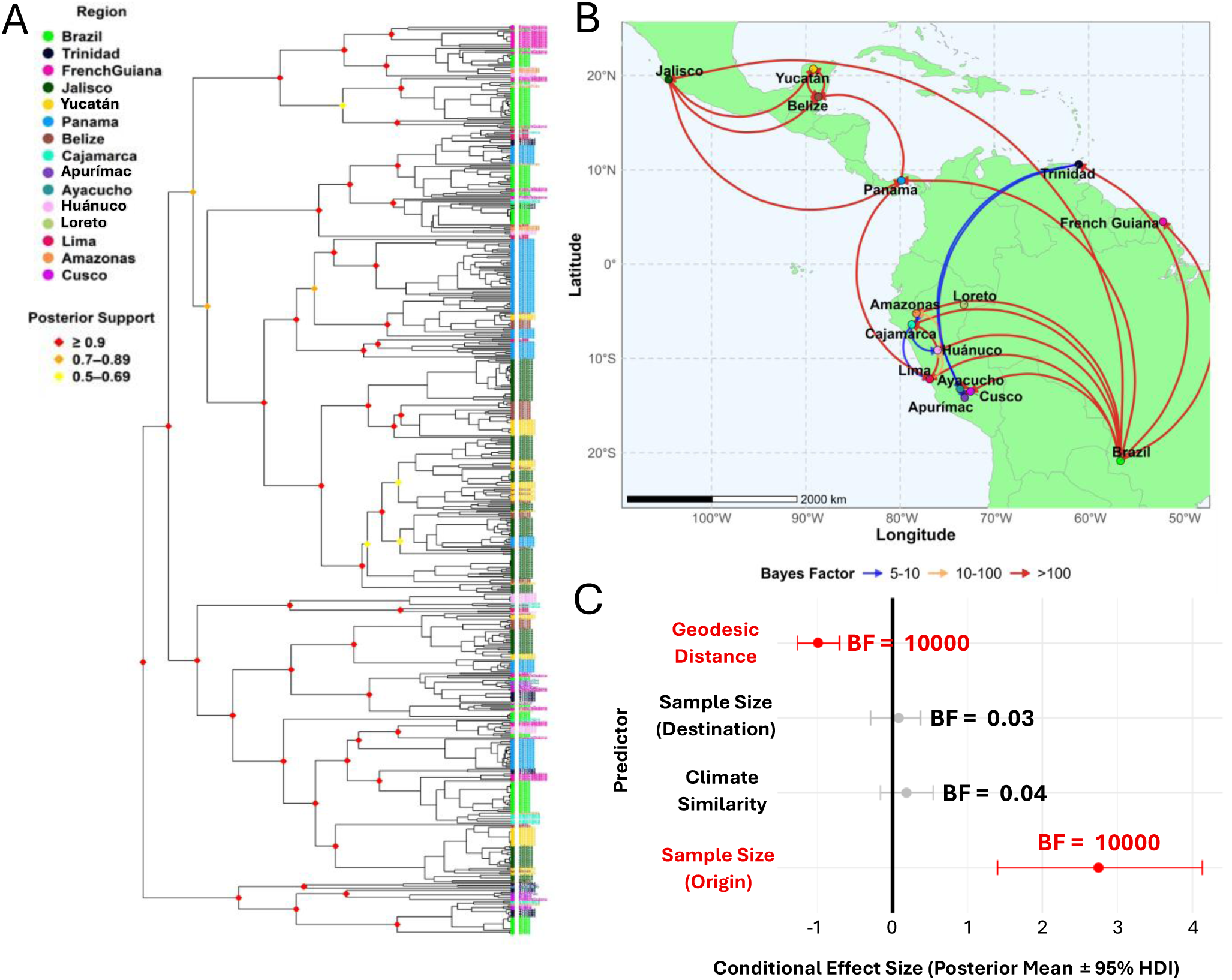
Bayesian phylogeographic reconstruction of DrBHV regional transitions across Latin America. Phylogeographic reconstruction of DrBHV regional transitions was done using a BSSVS analysis in BEAST, modelling region as a discrete trait. **(A)** Shows the inferred maximum clade credibility (MCC) tree. Tip points and labels were coloured by region, and coloured circles on deep internal nodes indicate posterior support categories. **(B)** Shows well-supported (BF > 5) regional transitions mapped onto Latin America. BF support was indicated by the colour of transition arrows. **(C)** Shows the conditional effect size plot (± 95% HDIs), i.e. the effect sizes when predictors were included in the model (δ = 1), for spatial and ecological predictors of recent (external) DrBHV regional transitions, derived from a BEAST GLM-CTMC analysis. Predictors with BF support > 5 were highlighted red. Sub-national regions belong to the following countries: Peru [Amazonas, Apurímac, Ayacucho, Cajamarca, Cusco, Huánuco, Lima, Loreto] and México [Jalisco, Yucatán].

Because terminal branches represent the most recent inferred spatial transitions in the phylogeny, transitions at the tips of the phylogeny provide the closest approximation to the contemporary inter-regional spread which might arise from a TV release under a field trial or deployment scenario. Therefore, following Faria et al. (2013) ^36^, we fitted a Generalised Linear Model-Continuous-Time Markov Chain (GLM-CTMC) analysis on external transitions. In this external-transition model, geodesic distance had a strong negative effect on inter-regional transition probability, indicating that recent DrBHV spread was more likely between geographically proximate regions (β = −0.995, HDI95: −1.268 to −0.708, inclusion probability = 1.000, BF = 10,000; Fig. 2C, Table S5). Inferred transition probability was also strongly associated with sampling effort in the origin region (β = 2.748, HDI95: 1.403 to 4.135, inclusion probability = 1.000, BF = 10,000; Fig 2C, Tables S5).

To test whether the same factors also shaped deeper historical spatial diffusion, we repeated the GLM-CTMC analysis using transitions that occurred on internal branches of the phylogenetic tree. This internal-transition model recovered the same negative effect of geodesic distance, although with weaker support than in the external model (β = −0.661, HDI95: −0.974 to −0.335, inclusion probability = 0.963, BF = 26.24; Table S5). Sampling effort at the origin was again strongly supported (β = 3.873, HDI95: 2.735 to 5.104, inclusion probability = 1.000, BF = 10,000; Table S5), as was sampling effort in the destination region (β = 0.833, HDI95: 0.436 to 1.237, inclusion probability = 0.997, BF = 319.57; Table S5). Together, these results suggest that geographic proximity has shaped DrBHV spatial diffusion across both recent and deeper evolutionary timescales, while also indicating that uneven regional sampling may influence inferred patterns of historical viral connectivity.

BSSVS identified 27 transitions between regions (BF > 5; Fig. 2B; Table S4). The most strongly supported BSSVS transitions (BF > 100) suggested a structured DrBHV metapopulation with Brazil, Jalisco (México), and Panamá acting as major sources of inter-regional lineages. Consistent with the GLM-CTMC results, many substantially supported inter-regional transitions occurred between geographically proximate regions, including Yucatán (México) and Belize, and among Peruvian regions (Fig. S1). We also identified substantially supported transitions from Brazil to Panamá, Brazil to Jalisco, and Panamá to Lima, consistent with effective viral connectivity between Central and South America, including routes spanning the Isthmus of Panamá (Fig. 2B; Table S4). The model inferred some additional longer-range connections from Trinidad and Tobago into southern Peru, but these transitions showed only moderate support (5 < BF < 10; Fig. S1; Table S4).

### Multiple Circulating DrBHV Strains with Varying Geographic Dissemination

Affinity propagation clustering of the MCC tree revealed 50 phylogenetic clusters, which we designated as DrBHV strains (Fig. S2). We found 79% of DrBHV-infected individuals harboured multiple DrBHV strains (range = 2 - 8, mean = 4.06 in co-infected bats). Strains were detected in 1–6 regions, and in 1–4 countries, highlighting variation in the geographic distribution of strains (Fig. 3(A); Table S6). Notably, 32 strains (64%) were detected in more than one region, indicating widespread circulation. However, the extent of strain sharing varied among regions, with some regions connected to multiple others and others showing more limited overlap in strain composition (Fig. 3(B)).

**Figure 3:**
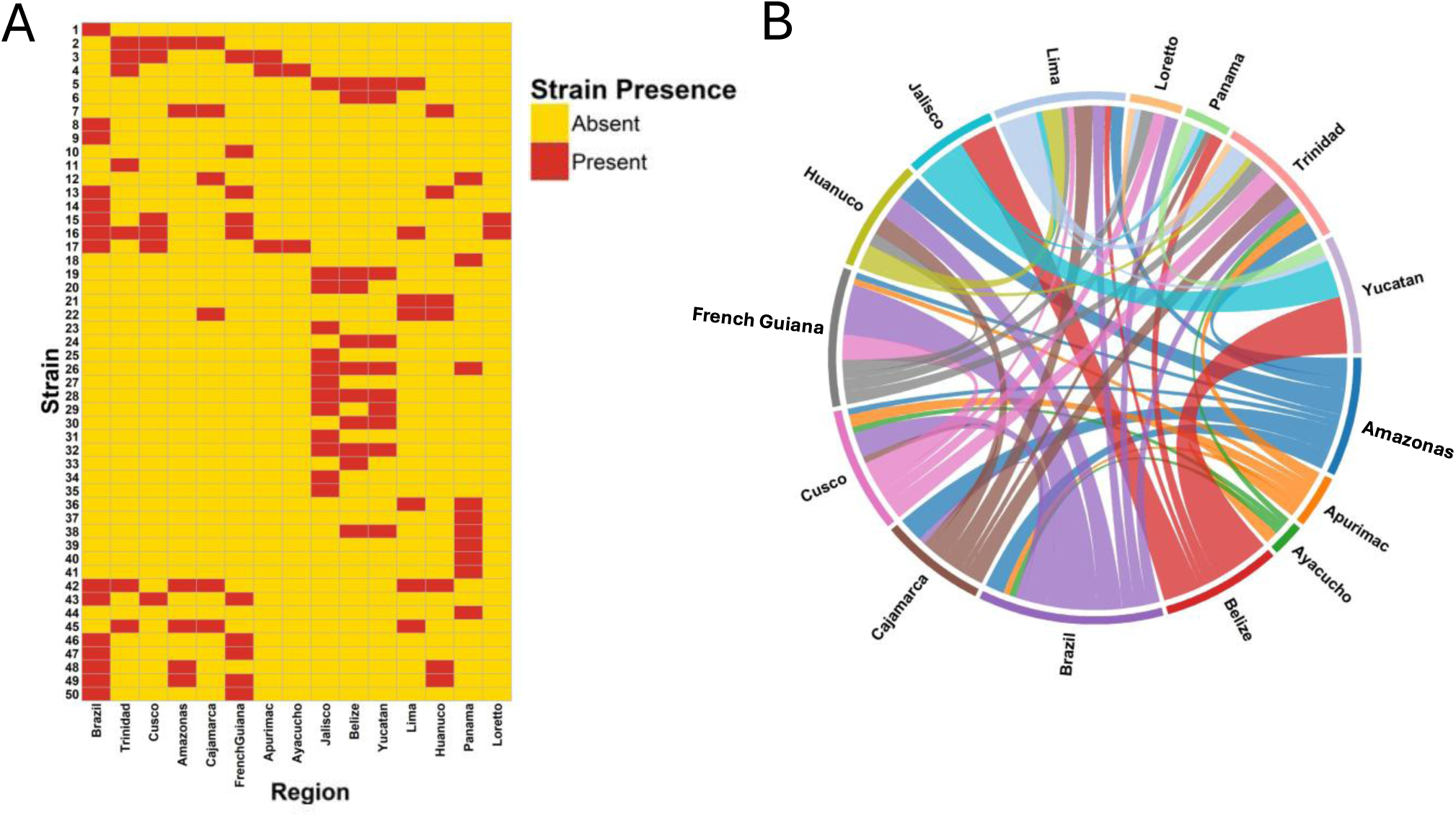
Geographic coverage and regional sharing of circulating DrBHV strains. **(A)** Shows a heatmap indicating strain presence or absence in each sampled region; and **(B)** Shows a chord plot visualising the quantity of strains shared between different regions. Thicker connections indicate higher quantities of strains shared. Sub-national regions belong to the following countries: Peru [Amazonas, Apurímac, Ayacucho, Cajamarca, Cusco, Huánuco, Lima, Loreto] and México [Jalisco, Yucatán].

### Genetic Similarity Does Not Constrain Co-Circulation and Co-infection of DrBHV Strains

To test whether strains are less likely to occur in regions containing closely-related strains, we fitted a binomial GLMM of strain presence/absence within each region as a function of (i) the minimum and mean genetic distance from the focal strain to strains already detected in that region, (ii) the minimum geodesic distance to the nearest region in which the focal strain was detected (a proxy for dispersal opportunity), (iii) regional strain richness, and (iv) sampling effort. Strain identity was included as a random intercept to account for baseline differences in overall prevalence among strains. Regional strain presence was strongly predicted by dispersal opportunity, with a lower probability of detection in regions farther from the nearest occupied region (β = −1.31, p < 0.0001, Fig. 4(C), Table S7) and increased with regional strain richness (β = 0.67, p = 0.001, Fig. 4(C), Table S7). Surprisingly, the mean genetic distance to regionally circulating strains was negatively associated with regional presence (β = −0.45, p = 0.040, Fig. 4(C), Table S7). This result indicates that strains were more likely to occur in regions where co-circulating strains were, on average, more genetically similar to the focal strain, likely reflecting geographic clustering of related strains. Thus, after controlling for dispersal opportunity and regional diversity, patterns of strain co-circulation were inconsistent with competitive exclusion between closely-related viruses and appear to instead be driven by ecological opportunity.

**Figure 4:**
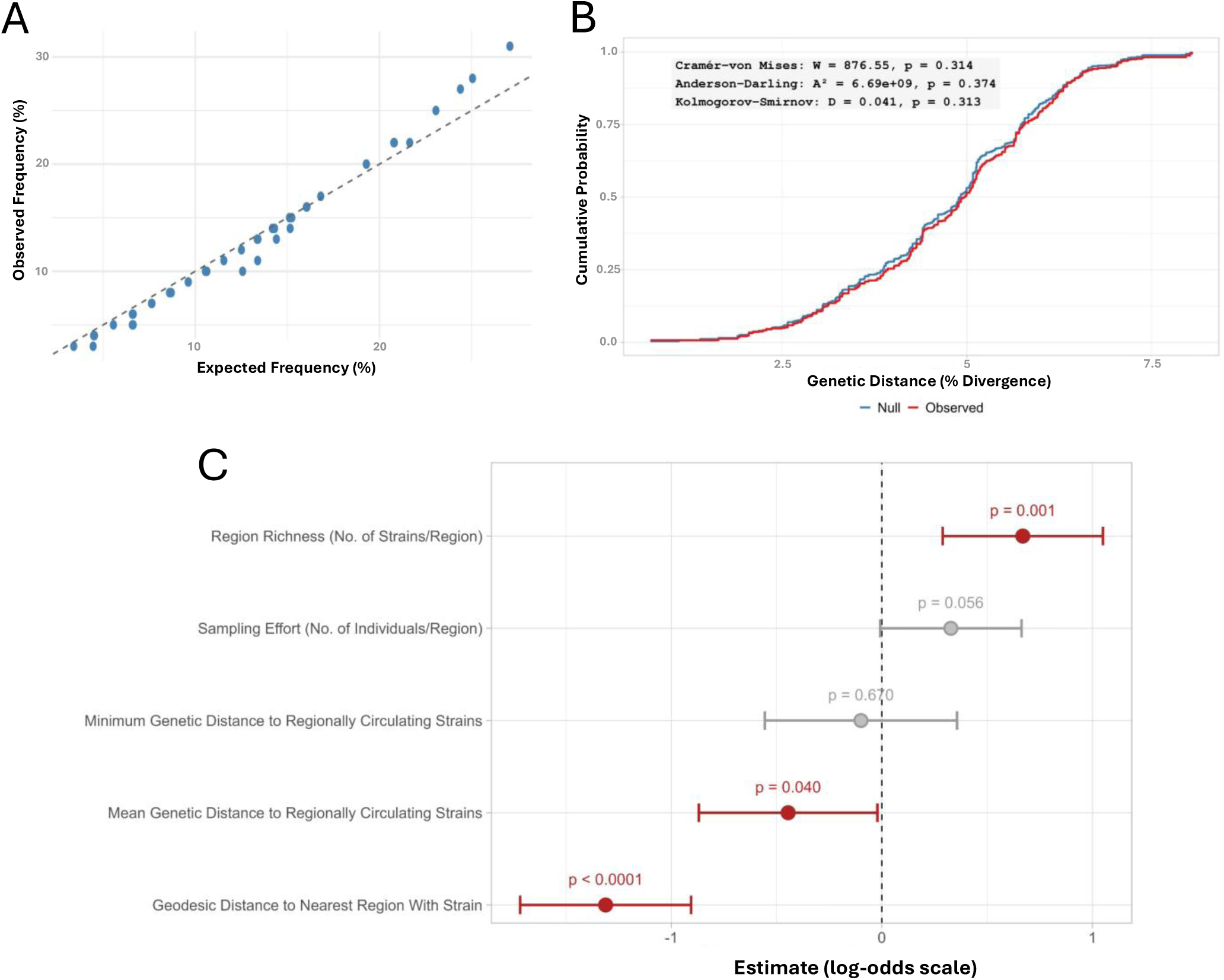
Regional co-circulation and within-host co-infection of DrBHV strains are not limited by genetic similarity. **(A)** Shows observed frequencies of strains in co-infected individuals compared to expected frequencies generated under a null model of random strain assignment, using 5,000 region-constrained, prevalence-weighted permutations. For each strain, we calculated a two-sided empirical permutation p-value based on the absolute deviation from the permutation mean. All strains were non-significant. **(B)** Shows empirical cumulative distribution functions (ECDFs) of the pooled distribution of observed pairwise genetic distances between co-infecting strains within individual bats (red line) compared to a null distribution generated from 5,000 region-constrained, prevalence-weighted permutations (blue line). Statistical differences between observed and null distributions were evaluated using non-parametric two-sample tests (Anderson–Darling, Cramér–von Mises, and Kolmogorov–Smirnov). **(C)** Shows fixed-effect estimates (± 95% CIs) from a binomial strain × region co-circulation GLMM predicting regional strain presence/absence from regional richness, sampling effort, minimum/mean genetic distance to strains detected in the region, and minimum geodesic distance to the nearest other region containing the focal strain (proxy for dispersal opportunity). Significant effects (Pr(>|z|) < 0.05) are highlighted in red.

To determine whether genetic similarity influences within-host co-infection beyond what is expected from local strain availability, we compared the pooled empirical cumulative distribution functions (ECDFs) of observed pairwise genetic distances between co-infecting strain pairs to those generated under a region-constrained, prevalence-weighted null model (5,000 permutations). Observed and null distance distributions were indistinguishable across the full range of genetic distances (Cramér–von Mises: W = 876.55, p = 0.314; Kolmogorov–Smirnov: D = 0.041, p = 0.313; Fig. 4(B)). Because very closely-related strains are particularly relevant to whether locally-derived, modified viral vectors could invade in the presence of co-circulating WT strains, we also evaluated differences at the lower tail. The minimum observed genetic divergence between co-infecting strains was 0.71%, which was identical to the minimum distance observed under the prevalence-weighted, region-constrained null. Further, an Anderson–Darling test, which places greater weight on distributional differences in the tails, found no evidence of deviation from null expectations (A² = 6.69 × 10⁹, p = 0.374; Fig. 4(B)), indicating that very closely-related strains naturally co-infect in proportion to their overall prevalence. Finally, we tested whether strains varied in their apparent ability to co-infect. Observed co-infection frequencies of individual strains were indistinguishable from a null model which accounted for regional coverage and local prevalence (empirical permutation test: p > 0.05, 5,000 iterations), implying broadly similar capacity for co-infection among strains (Fig. 4(A)). Alongside the regional analysis, these results argue against strong competition or cross-immunity among closely-related DrBHV strains and suggest that effective vectors will not require exceptional superinfection capability, removing major hypothesised barriers to strain selection for vaccine vectors.

## Discussion

TVs offer a theoretically promising alternative to bat culling for rabies management in Latin America, but their operational feasibility hinges on identifying a safe, host-restricted viral vector that is likely to spread predictably through bat populations while evading potential interference from the endemic WT strains. Here, we show that DrBHV comprises a single, well-supported clade across the broad, geographic range of vampire bats, with no detectable regional differences in prevalence. Contemporary spatial spread was predicted by geographic proximity, implying that dissemination is most likely to proceed via stepwise diffusion through neighbouring regions rather than frequent long-distance jumps. Finally, we found strains co-infected in proportion to their local prevalence and no evidence that genetic similarity systematically constrained within-host co-infection or regional coexistence. These findings strengthen the case for DrBHV as a vector for a transmissible rabies vaccine and demonstrate how interrogating patterns of WT infection in natural populations can anticipate future vaccine dissemination routes, guide vector selection for vaccine design, and inform governance and monitoring strategies.

Our phylogenetic analysis demonstrated that DrBHV occurs across the range of vampire bats, forming a single, strongly-supported monophyletic clade which spanned major geographic barriers (e.g. the Andes mountains) and a Caribbean island population. This range-wide continuity supports a long-standing association between DrBHV and vampire bats and aligns with current interpretations that mammalian BHVs co-evolved with their hosts rather than frequently switching into novel host species ^37^. For the TV concept, this ancient association is encouraging because it suggests a stable evolutionary relationship with a lower likelihood of unanticipated evolutionary changes following release (e.g., shifts in pathogenicity or host range). Although our study was not intended to detect divergent BHVs in vampire bats or to detect DrBHV in non-vampire bat species, these represent additional dimensions of host-specificity which will need to be evaluated to fully understand the extent to which DrBHV-based TVs might affect non-target hosts ^21,27^. We also noted substantial, albeit not statistically significant, variation in prevalence across the seven countries analysed here. It is currently unclear whether this variation represents genuine differences in prevalence which could not be detected due to the low sample sizes in some geographic areas, or that prevalence was similar across regions with variation in primer performance for locally-circulating strains creating apparent differences, an issue previously documented for Zika virus and SARS-CoV-2 assays ^38,39^. Strain-agnostic diagnostics, such as pan-BHV PCRs, would be a promising alternative to generate less-biased estimates of prevalence. Additionally, the low prevalence observed in French Guiana, where the samples were more than 10 years old, may partly reflect DNA degradation, which can be particularly problematic when amplifying larger amplicons such as those targeted here ^40,41^. If confirmed to be biologically meaningful, regional variation in prevalence would suggest that achieving management targets with a DrBHV-vectored vaccine may require substantially greater deployment effort in countries where baseline prevalence is consistently or periodically low. We also found that reproductively active individuals had 62% lower odds of infection, suggesting that seasonal shifts in breeding phenology and sampling timing could influence observed prevalence patterns and short-term TV dissemination conditions ^42^.

Because TVs are designed to circulate autonomously in wildlife, their benefits and risks are not confined to the release site. Host movement and transmission networks can carry a vaccine strain across administrative boundaries. Regional governance is therefore central to responsible use, requiring shared oversight, transparent monitoring for unintended spread, and pre-agreed escalation criteria to prevent unmanaged cross-border risks ^27^. Our phylogeographic analysis provides an empirical basis for anticipating how a DrBHV-vectored TV may spread, demonstrating that the probability of inter-regional spread strongly decreased with geographic distance, particularly when considering a proxy for recent spread. This suggests that early post-release dissemination would be most likely to occur between neighbouring regions, rather than through unpredictable long-distance jumps. Monitoring of a released TV could therefore prioritise areas closest to the release site, making surveillance more targeted, feasible, and rapidly actionable. This distance-constrained pattern is encouraging from a governance and logistics perspective, because it implies that cross-border risks may be manageable through targeted regional surveillance networks rather than requiring immediate continent-wide monitoring following release.

Despite this overarching distance-constrained pattern across both recent and historical spatial diffusion, the BSSVS reconstruction also identified a small number of connections between geographically distant regions. If verified, such unpredictable, long-range cross-jurisdictional spread would introduce a major governance challenge. However, given that vampire bats have small home ranges and are non-migratory, we speculate that apparent long-range jumps are more likely to reflect ancient diversification of DrBHV strains or multi-step diffusion across poorly sampled or unsampled regions, rather than discrete long-distance dispersal events directly relevant to TV deployment ^42,43^. Although it would have been ideal to include a time-scaled phylogeographic analysis to estimate how quickly these inter-regional transitions occur, this was not supported by the available data because of limited temporal signal. Moreover, because Betaherpesviruses establish latent infections and can later reactivate, sampling dates may not correspond closely to the timing of primary infection, meaning that some apparent “transitions” could instead reflect reactivation of long-standing latent infections. Importantly, TVs are expected to remove or silence their transgenes, resulting in an effective reversion to the WT ^35^. Temporal analyses of spread, together with estimates of TV evolutionary stability, could determine whether reversion would be expected to pre-date cross-boundary spread, thereby reducing regulatory concerns.

Improving strain resolution is important for TV development because knowledge of which strains circulate, and where, is fundamental for selecting an appropriate vector backbone and anticipating spatial differences in vector suitability, spread, and long-term persistence. Consistent with this, increasing genomic resolution substantially expanded inferred diversity relative to previous studies. We identified 50 circulating strains across the study region, including 18 in Peru alone, compared with 11 strains previously reported in Peru ^30^. This suggests that lower-resolution assays may underestimate the extent of standing viral diversity relevant to vector selection and deployment planning. This standing diversity is important not only for selecting an appropriate vaccine backbone, but also because any candidate TV would be released into host populations already containing multiple endemic WT lineages.

A DrBHV-based TV construct would need to superinfect hosts that already carry WT DrBHV strains, and then persist and transmit in the presence of those strains. Our co-infection and co-circulation analyses provide three lines of evidence that WT strain diversity is unlikely to impede the establishment and transmission of DrBHV-based TVs. First, although strains could plausibly differ in traits that influence superinfection (e.g., growth kinetics, immune evasion, or superinfection exclusion ^44–47^), we found no evidence that particular DrBHV strains were consistently over- or under-represented among co-infected hosts. This pattern is encouraging, because it reduces the likelihood that TV success would require identifying and sourcing rare “high-performance” backbones. Second, we tested whether co-infections are preferentially composed of closely-versus distantly-related strains using gB sequences, which encodes an antigenic entry glycoprotein and is therefore a plausible locus for immune-mediated interference. Strong cross-immunity could blunt TV impact and necessitate more intensive or repeated deployment ^23,29^. However, the genetic distances between co-infecting strains matched null expectations across the full distribution, including among the most closely-related pairs, indicating that genetically similar strains can co-infect and, more generally, strains co-infect in proportion to their local prevalence. While cross-neutralisation assays are needed to quantify the strength of cross-protection directly, these patterns argue against strong, consistent exclusion at gB that would systematically prevent co-infection by closely-related DrBHV lineages. Third, at the regional scale, strain co-circulation was better explained by dispersal opportunity and local strain richness than by competitive sorting. These patterns are consistent with shared ancestry and spatially-structured diffusion, rather than competitive exclusion among closely-related strains. This diffusion-based interpretation aligns with prior work in Peru, which also favoured dispersal and local emergence over competition as drivers of strain ranges ^30^. Taken together, these results suggest that circulation of related WT strains is unlikely to block vaccine invasion, supporting the feasibility of using endemic, locally circulating DrBHV backbones. This is operationally and ethically important since importing viral backbones between regions could introduce novel DrBHV diversity into naïve areas – raising biosafety, ecological, and regulatory concerns.

An important caveat to our conclusions on superinfection is that our analyses address whether co-infection occurs, not whether superinfection is equally productive. Even if WT strains do not prevent infection outright, more nuanced effects (e.g., viral titre, transmission, tissue tropism, or transgene expression) could influence TV performance and should be tested experimentally (e.g., cross-neutralisation, controlled co-infection/challenge studies, and assays of shedding and transmission). We also could not resolve homologous (same-strain) reinfection because our strain definitions are cluster-based and our co-infection framework evaluates pairs of distinct strains. However, same strain co-infection underpins whether a locally-derived TV backbone can infect hosts already carrying that same strain. Distinguishing true homologous reinfection from latency/reactivation or within-host evolution will likely require longitudinal designs and controlled *in vivo* experiments using genetically-barcoded viruses that can separate reinfection from recrudescence.

Overall, this study helps move TVs from proof-of-concept toward deployable interventions by linking the ecology of viral vectors to the practical constraints of design, deployment, and governance. For design, our findings suggest that DrBHV-based TVs are unlikely to face systematic interference from endemic strain diversity, supporting the feasibility of using local, endemic backbones as opposed to importing viruses between regions – an important biosafety and regulatory advantage. Just as importantly, detecting DrBHV in every sampled country indicates that suitable vectors are likely to be available and effective across most rabies-endemic regions of the *D. rotundus* range. For deployment, the proximity-driven structure of spread implies that dissemination can be planned and evaluated as a stepwise process through connected host networks, enabling rational choices about where to seed vaccines to maximise reach and where to start with more contained, monitorable trials. Finally, because self-disseminating interventions create cross-border externalities by design, our results underscore the need for regional governance frameworks that couple staged releases with transparent, shared surveillance and pre-agreed triggers for escalation or stopping, so that technical performance gains translate into responsible, scalable public-health benefit.

## Materials and Methods

### Samples, Study Region and Collaborative Approaches

#### Samples and Study Region

We analysed 229 oropharyngeal swabs from wild *D. rotundus* captured between 2011 and 2024 in 15 regions across seven Latin American and Caribbean countries, spanning most of the species’ native range (Fig. 1(A)). Regions were defined at the country level unless multiple first-level administrative subdivisions were sampled within a country, in which case each subdivision was treated as a separate region. Regions sampled (n = number of samples) were: eight regions in Peru [Lima (n = 20), Apurímac (n = 9), Ayacucho (n = 20), Cusco (n = 10), Huánuco (n = 13), Cajamarca (n = 12), Amazonas (n = 4) and Loreto (n = 6)]; one state in Brazil (Mato Grosso do Sul, n = 26); two states in México (Jalisco, n = 19; Yucatán, n = 20); Orange Walk District in Belize (n = 19); Panamá Oeste province in Panamá (n = 13); Roura commune in French Guiana (n = 25); and Sangre Grande region in Trinidad & Tobago (n = 13). Analyses were conducted at the regional level to maximise spatial resolution while maintaining adequate sample sizes. Some regions contained multiple sites, which were pooled because of the large geographic scale of the study (Table 1). Swabs were stored in 1 mL RNALater (#AM7021, ThermoFisher Scientific) or 1 mL DNA/RNA Shield (#R1100-250, Zymo Research).

#### Location of Work and Division of Labour

Peruvian samples were collected during previous field expeditions and subsequently extracted and tested at the Medical Research Council–University of Glasgow Centre for Virus Research (CVR). Brazilian and Mexican samples were collected locally and then stored at -80°C in specific institutes (Brazil: Centro Estadual de Vigilância em Saúde; México: U.S. Geological Survey – National Wildlife Health Centre [USGS-NWHC]) before shipment to the CVR for extraction and testing. Samples from French Guiana, Belize and Panamá had been extracted and stored at specific institutes (French Guiana: Institut Pasteur de la Guyane; Panamá/Belize: University of Oklahoma), and were shipped to the CVR for testing. Trinidad & Tobago samples were collected, extracted and tested locally (University of the West Indies); positive PCR products were then sent to the CVR. All selected samples were sequenced at the CVR.

#### Research and Animal Ethics Approval

Peruvian sample collection was authorised by the Peruvian government (RD-009-2015-SERFOR-DGGSPFFS, RD-264-2015-SERFOR-DGGSPFFS, RD-142-2015-SERFOR-DGGSPFFS, RD-054-2016-SERFOR-DGGSPFFS), and animal capture, handling and sampling protocols were approved by the University of Glasgow School of Medical, Veterinary and Life Sciences Research Ethics Committee (EA59/23). Mexican samples were collected under SEMARNAT permit 19/LW-0120/11/20, with capture, handling and sampling protocols approved by the USGS-NWHC Animal Care and Use Committee (EP190528.A1). Brazilian sample collection was authorised by the Ministry of the Environment (SISBIO, proc. no. 10615-8), and protocols were approved by the Animal Use Ethics Committee at the Federal University of Mato Grosso do Sul (1244/2022). Trinidadian samples were collected under Ministry of Agriculture, Land and Fisheries Wildlife Section permit 2021/11/22-094, with protocols approved by The University of the West Indies Research Ethics Committee (CREC-SA.2296/08/2023). Panamanian samples were collected under Ministerio de Ambiente (MiAmbiente) permit SE/A-26-2020, with animal procedures approved by the Smithsonian Tropical Research Institute (STRI) Animal Care and Use Committee (SI-21034). Belizean samples were collected under Belize Forest Department permit FD/WL/1/19(09), with protocols approved by the American Museum of Natural History Institutional Animal Care and Use Committee (AMNHIACUC-20190129). French Guianan samples were collected under Regional Scientific Council for Natural Heritage (CSRPN) authorisation 59; ethics permits were not required because bats are not protected in French Guiana.

### Sample Preparation and Sequencing Pipeline

#### Nucleic Acid Extraction

Samples extracted at the CVR followed buffer-specific nucleic acid extraction protocols. Swabs stored in RNALater were inactivated with Buffer RLT (#79216, Qiagen) in a derogated CL3 facility using a previously described double swab-lysis protocol ^28^, then extracted on a KingFisher Flex 96 automated platform (ThermoFisher Scientific) with the BioSprint One-For-All Vet Kit (#947057, Qiagen) following a protocol optimised for detecting viral nucleic acid ^28^. Swabs stored in DNA/RNA Shield were extracted either on the KingFisher Flex 96 using the Quick-DNA/RNA High-Throughput Kit (#R2150, Zymo Research) according to the manufacturer’s protocol, or with the Quick-DNA/RNA Viral Spin Column Kit (#R7021, Zymo Research).

#### Amplification of the DrBHV DPOL-gB Region

Primers were derived from a previous PrimalScheme panel designed to amplify a 12 kbp region of the DrBHV genome ^30,48^. The outermost primer pair amplified poorly in pilot Peruvian samples, likely because the 12 kbp target exceeded the polymerase’s reliable range ^49^. We therefore selected a primer pair flanking a 3,919 bp region containing 79 bp of UL56 at the 3′ end, the full 2,991 bp glycoprotein B (gB; UL55) gene and 849 bp of DNA polymerase (DPOL; UL54) at the 5′ end (F: DrBHVprimer_3_LEFT, 5′-GAAGGCCATGCACTTGTACACC-3′; R: DrBHVprimer_4_RIGHT, 5′-AACGAGGACGACGACGATTTTG-3′). We targeted gB because of its hypervariability and extensive use in cytomegalovirus (CMV) strain-typing studies in humans and animal models ^50–54^. For each sample, 2.5 µL of extract was amplified with Q5 high-fidelity polymerase (#M0491S, New England Biolabs) in 5X reaction buffer. Cycling conditions were: 98 °C for 30 s; 45 cycles of 98 °C for 15 s, 65 °C for 30 s and 72 °C for 3.5 min; then 72 °C for 10 min and a 4 °C hold. Successful amplification was confirmed by a ∼4 kbp band on a 0.9% agarose gel.

#### Library Preparation and Sequencing of DPOL-gB Amplicons

Libraries were prepared from 200 fmol of Ampure XP (#A63881, Beckman Coulter)–cleaned PCR product using the Native Barcoding Kit 96 V14 (#SQK-NBD114.96, Oxford Nanopore Technology), following the manufacturer’s instructions with minor modifications. DNA end-preparation and barcode ligation were combined ^55^, and Ampure XP bead clean-ups were performed on individual samples after barcode ligation instead of heat/EDTA inactivation. DNA concentration and quality were assessed with the Qubit dsDNA High Sensitivity assay (#Q32854, ThermoFisher Scientific). Multiplexed libraries were sequenced for 40–48 h on a MinION Mk1B device (#MIN-101B, Oxford Nanopore Technology) with R10.4.1 flow cells (#FLO-MIN114, Oxford Nanopore Technology).

### Bioinformatic Processing of Raw Reads

After sequencing, .pod5 files were basecalled on a high-performance computing server using Dorado v0.9.6 with the SUP (super high accuracy) model. FastQC v0.12.1 was used for initial quality control. Raw reads were (1) quality-filtered to remove reads with Q < 20 (c = 0.99) using nanoq v0.10.0, (2) length-filtered to retain reads between 3.5 and 4.5 kbp using seqkit v2.8.2, and (3) demultiplexed with Guppy v6.5.7. For each barcode (sample), filtered reads were mapped to a reference DrBHV DPOL-gB sequence from a previously annotated genome, 10148_c5 (SRR11789719) ^56^, using Minimap2 v2.28. Mapped reads were extracted (and reverse-complemented when required) with Samtools v1.22 and converted to FASTA format using seqkit v2.8.2. To correct for lower sequencing accuracy and systematic errors in ONT data, FASTA sequences from each barcoded pool were clustered with CD-HIT v4.8.1 (cd-hit-est, 95% identity, c = 0.95). Clusters with <10 reads were discarded. Reads in remaining clusters were extracted and aligned with MAFFT v7.475, and consensus sequences were generated using an in-house Python script, calling bases with frequency >50% and assigning N otherwise.

### Determining Regional and Host-related Factors Affecting DrBHV Prevalence

To identify regional and host-related predictors of DrBHV prevalence, we fitted binomial GLMMs in R using *glmmTMB*, with DrBHV infection status as the binary response. Candidate fixed effects included region (Lima, Apurímac, Ayacucho, Cusco, Huánuco, Cajamarca, Amazonas, Loreto, Brazil, Jalisco, Yucatán, Belize and Panamá), sex (male/female), age class (adult/juvenile/sub-adult), reproductive status (yes/no), weight, and forearm length, with capture year included as a random intercept. Samples from Trinidad & Tobago and French Guiana were excluded because of missing metadata. Prior to model selection, we assessed collinearity among candidate predictors using variation inflation factors (VIFs) and pairwise correlations. Forearm length showed the highest variance inflation factor (VIF = 3.67), and forearm length and weight were also moderately positively correlated (Pearson’s r = 0.593, 95% CI: 0.491–0.678, p < 0.001). Because both variables capture overlapping aspects of body size, we constrained the candidate model sets so that only one of these terms could be included in any given model. We then ranked the resulting constrained candidate set using *MuMIn::dredge* and corrected Akaike’s Information Criterion (AICc) in R, and drew inference from the best-supported model.

Because female reproductive status could not be partitioned into finer categories (e.g. pregnant or lactating) without substantial loss of statistical power, we fitted an additional binomial GLMM including a sex × reproductive status interaction and compared it with the best-supported additive model using AICc and a chi-squared likelihood ratio test (LRT).

### Phylogenetic Analyses

#### Assessing DrBHV Monophyly Across the Study Region

To obtain homologous gB-DPOL sequences from other mammalian Betaherpesviruses, we used BLASTn to search the *core_nt* database with random *D. rotundus*–derived sequences from this study. We selected 29 annotated entries spanning six mammalian groups known to host Betaherpesviruses - humans, non-human primates, rodents, bats, shrews and cavies ^57–62^ - and extracted the homologous gB-DPOL regions (Table S3). These sequences were aligned with all *D. rotundus* sequences using MAFFT v7.475, manually inspected and trimmed in AliView v1.28 to remove sites outside the primer binding regions, and further processed in TrimAl v1.5.0 (using the Maximum Likelihood (ML) optimised *-automated1* option) to remove uninformative gappy positions arising due to large sequence divergence. ML phylogenies were inferred from the trimmed alignment in IQ-TREE 2 v2.4.0 under a GTR+F+R10 substitution model selected by ModelFinder (lowest Bayesian Information Criterion, BIC) with 10,000 ultrafast bootstrap (UFBoot) replicates. Resulting trees were processed and visualised in R using *ape, ggtree* and *ggplot2*. The tree was rooted using KC465951.1_HHV-6A_GS (Human herpesvirus 6A, Roseolovirus) as an outgroup, although the root was removed for final visualisation using *ape*. Animal silhouettes were added for visual clarity using PhyloPic v2.0. Pairwise genetic distances between sequences were computed as uncorrected p-distances (model = "raw") from the alignment, with pairwise deletion of sites containing gaps, using *ape*.

#### Regional Transitions of DrBHV Across Latin America and Strain Delineation

All DrBHV sequences were aligned in MAFFT v7.475 using the L-INS-i algorithm for increased accuracy. Alignments were manually inspected and trimmed in AliView v1.28 to remove primer binding regions. Bayesian phylogenetic inference was performed in BEAST v1.10.4 ^63^ for 2 × 10^8^ generations, sampling every 1 × 10^4^ steps, under a strict clock (rate = 1.0, reflecting limited longitudinal data and possible non-correspondence of sampling and infection dates due to viral latency), a constant population size coalescent prior and a TIM+F+I+G4 substitution model (lowest BIC in IQ-TREE 2). All other priors were set to default. The TIM+F+I+G4 model was coded into the BEAUTI XML file. Sequences were assigned a region trait, modelled as a separate partition under a discrete phylogeographic model with an asymmetric BSSVS procedure to infer ancestral states and regional transitions. Outputs were assessed in Tracer v1.7.2 (burn-in: 10%; 2 × 10^7^ states; all ESS > 200). A MCC tree was generated in TreeAnnotator v1.10.4 and visualised in R using *treeio, tidytree*, *phytools* and *ape*. Strains were delineated from the MCC tree with the *apcluster* package in R, as previously used for DrBHV strain delineation based on a shorter gB fragment ^30^. Regional strain presence and strain sharing among regions (chord plots) were summarised in R using *ComplexHeatmap* and *circlize*, respectively. The relative support for individual regional transitions was assessed by calculating Bayes Factors (BFs) from the BSSVS procedure in BEAST, as previously described ^43,64^. MCMC chains were processed in R using the *tidyverse* package suite. Briefly, for each possible transition between regions, a binary indicator variable δ was estimated, where δ = 1 indicates that a transition is included in the model and δ = 0 indicates exclusion. Posterior inclusion probabilities (p_ppp_) for each transition were obtained as the mean of the indicator variables after removing a 10% burn-in.

The prior inclusion probability (q_ppp_) for each transition was calculated using a Poisson prior with mean λ = 14 on the expected number of non-zero transition rates among all K = N × (N-1) possible directional transitions between N = 15 regions, such that q_ppp_ = λ / K. Bayes factors (BFs) were then calculated as the ratio of posterior to prior odds of inclusion:

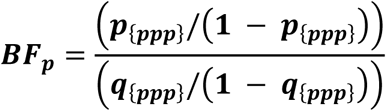

where larger values indicate stronger support for a transition than expected under the prior. We considered transitions with BF > 5 to be well supported and classified BF support as 5–10 (moderate), 10–100 (strong) and >100 (decisive). Transitions with infinite posterior odds (i.e., p_ppp_ = 1) had BF values capped to 10,000. BFs were cross-checked against SpreaD3 output ^65^. Using latitude–longitude metadata for sampling sites, significant transitions were mapped across Latin America using *ggplot2, sf, ggrepel* and *ggspatial* in R.

#### Determining Predictors of Inter-Regional DrBHV Transitions

To determine predictors of DrBHV regional transitions, we ran a separate BEAST analysis modelling the region trait using a GLM-CTMC framework with the following settings: 1 × 10^8^ generations with sampling every 5 × 10^3^ steps, strict clock, constant population size coalescent model, default priors and the same TIM+F+I+G4 substitution model. To distinguish predictors of recent spatial spread from those associated with deeper historical diffusion, we fitted separate GLM-CTMC models to external and internal transitions, following Faria et al. (2013) ^36^. External transitions were defined as transitions occurring on terminal branches and were treated as the primary analysis because they provide the closest approximation to contemporary inter-regional spread relevant to a vaccine release or field-trial scenario. Internal transitions were analysed separately to test whether the same predictors also explained deeper historical spatial diffusion across the phylogeny. We tested three predictors in both the external- and internal-transition models:

*(1) Geodesic distance:* To test whether DrBHV inter-regional transmission can be predicted simply by geographic proximity. This was calculated as the geographic distance (utilising latitude and longitude coordinates) between regions using the Haversine formula, which accounts for Earth’s curvature, implemented in R with the geosphere package. Distances were expressed as great-circle kilometres, and a symmetric pairwise distance matrix was produced for all regional combinations.
*(2) Climate Similarity:* To test whether inferred DrBHV transitions between regions reflect ecological connectivity (shared environmental suitability) rather than geographic proximity alone, we quantified climatic similarity among regions. Environmental covariates are routinely included in phylogeographic GLM frameworks, and are often supported as significant predictors of transition rates in empirical applications across multiple viral systems ^66–69^. This is further motivated by evidence that climate shapes the distribution of vampire bats and the spillover risk of VBRV across the Americas ^33,34,70^. We extracted 19 bioclimatic variables from the WorldClim v2 dataset at 10-arcminute (∼18 km) resolution for each site using the geodata and raster packages in R. Bioclimatic values were standardised (mean = 0, SD = 1) to account for differences in variable magnitude, and Euclidean distances between region coordinates were calculated in this multivariate climate space. These distances were then scaled to a 0–1 climate similarity index, where 1 represents the most climatically similar pair of regions and 0 represents the most dissimilar, forming a symmetric pairwise similarity matrix.
*(3) Sampling Effort:* Total number of samples per region, included as both origin and destination predictors to test whether differences in sampling intensity affected inferred transitions to and from regions.

As above, appropriate burn-in and model convergence were assessed in Tracer v-1.7.2 (ESS > 200; burn-in = 10% states). We calculated predictor support using BFs as described above, but with a different prior inclusion probability (q_ppp_). A Bernoulli distribution was used for δ, meaning there was an equal probability of inclusion or exclusion for each variable such that q_ppp_ = 0.5 ^71^. Predictors with infinite posterior odds (i.e., p_ppp_ = 1) had BF values capped to 10,000. Well-supported transitions were determined when BF > 5. An effect size plot was generated using *ggplot2* in R.

### Assessing Genetic Constraints on Co-Circulation and Co-infection of DrBHV Strains

We evaluated whether genetic similarity constrains co-circulation and co-infection of closely-related DrBHV strains by testing (i) whether genetically similar strains are less likely to co-infect individual bats than expected given local strain availability, and (ii) whether genetically similar strains are less likely to co-circulate within the same region after accounting for dispersal opportunity and sampling intensity. All analyses were conducted at the regional scale (rather than country) to ensure that availability pools reflect local circulation patterns and to avoid bias where sampling sites may be geographically closer across national borders than within them. We used one representative sequence per strain for all genetic distance calculations to avoid redundancy.

To test whether genetically similar strains were more or less likely to co-infect within individuals, we calculated pairwise genetic distances between all strain representatives directly from the aligned nucleotide sequences as raw p-distances (proportion of sites differing) using *dist.dna()* in *ape*. For each multi-strain–infected bat, we enumerated all unique pairs of co-infecting strains (individuals with only one detected strain were excluded) and retrieved the corresponding observed pairwise genetic distances from the alignment-based distance matrix. Null expectations were generated using a prevalence-weighted, region-constrained sampling procedure that preserves local availability. For each observed co-infecting pair, we generated 5,000 null pairs by sampling two distinct strains from the same region as the observed pair, with sampling probabilities proportional to each strain’s observed regional prevalence (based on all infections, not only co-infections). Identical strain draws were rejected. Genetic distances for sampled null pairs were retrieved from the same alignment-based distance matrix and pooled across observed pairs to form the overall null distribution. Observed and null distance distributions were compared using empirical cumulative distribution functions (ECDFs) and non-parametric two-sample tests (Cramér–von Mises, Kolmogorov–Smirnov, and Anderson–Darling ^72^) implemented in the *twosamples* package. Significance was defined when p < 0.05.

To test whether genetic similarity or dispersal opportunity predicts whether a strain occurs in a given region, we constructed a strain × region presence/absence dataset. For each focal strain and region, we calculated (i) the minimum and mean genetic distance from the focal strain to all other strains detected in that region (excluding the focal strain when present), using the same representative sequences and the previously generated alignment-based distance matrix; and (ii) the minimum geodesic distance (Haversine) from the focal region to the nearest other region in which the focal strain was detected, using latitude/longitude coordinates and the *geosphere* package. Strains observed in only a single region were excluded because the nearest “other occupied region” distance was undefined. We additionally quantified regional sampling effort (number of unique individuals sampled per region) and regional strain richness (number of strains detected per region). All fixed effects were z-scaled. Collinearity diagnostics using the *check_collinearity()* function in *performance* revealed all VIFs were < 2.8. Strain presence/absence across regions was modelled using a binomial GLMM implemented in *glmmTMB* with fixed effects for: (i) minimum and mean genetic distance to locally circulating strains, (ii) minimum geodesic distance to (other) region with the strain, (iii) regional strain richness, and (iv) sampling effort covariates. Strain identity was included as a random intercept to account for baseline differences in overall prevalence among strains. Significance was defined when Pr(>|z|) < 0.05.

Finally, we tested whether any DrBHV strains were disproportionately represented among multi-strain–infected individuals after accounting for regional strain availability and local prevalence. We restricted this analysis to bats harbouring more than one unique strain and implemented a prevalence-weighted, region-constrained permutation test (5,000 iterations) in which, for each individual, we randomly sampled, without replacement, the same number of strains observed in that individual from the corresponding regional pool, with sampling probabilities reflecting regional strain prevalence.

This preserved each individual’s observed co-infection load and regional availability while breaking any non-random association between individuals and particular strains. Observed strain frequencies among co-infected bats were compared to the permutation null distribution, and for each strain we computed a two-sided empirical permutation p-value based on the absolute deviation from the permutation mean (p < 0.05 considered significant). Expected frequencies were summarised as permutation means and visualised alongside observed frequencies using an observed-versus-expected scatterplot with a 1:1 reference line using *ggplot2*.

## Acknowledgements

D.G.S and H.M were supported by a Wellcome Senior Research Fellowship (217221/Z/19/Z). H.M was also supported by a Henry Dryerre PhD Scholarship (PHD010657) issued by The Carnegie Trust for the Universities of Scotland. D.G.S, L.B. and A.B. were supported by the NSF/BBSRC Ecology and Evolution of Infectious Diseases program (DEB 2011069; BB/V003798/1). D.G.S was also supported by a Philip Leverhulme Prize from the Leverhulme Trust (PLP-2020-362). L.B. was also supported by a Natural Environment Research Council (NERC) award (NE/X011747/1). S.S. and G.G.C. are supported by grants from the National Science Foundation (DEB-2508535, DEB-2508536). A.R., C.C., L.R. and N.M. were supported by a Wellcome Trust Grant (226142/Z/22/Z). D.J.B was supported by the National Geographic Society (NGS-55503R-19) and Edward Mallinckrodt, Jr. Foundation. M.C.S. was supported by an appointment to the Intelligence Community Postdoctoral Research Fellowship Program at University of Oklahoma administered by Oak Ridge Institute for Science and Education through an interagency agreement between the U.S. Department of Energy and the Office of the Director of National Intelligence. E.F. was supported by a Research Grant (306345/2019-6) from the National Council for Scientific and Technological Development. Additional funding was provided by the UK Medical Research Council (MRC) through core support to the MRC-University of Glasgow Centre for Virus Research (Virus Cross Species Transmission Programme: MC\UU\00034/3). We would like to thank Fernando Gonçalves and Mauro Galetti for their help in coordinating internal sample shipments across Brazil. We would also like to thank Ana Da Silva Felipe, Katherine Smollett, Daniel Mair and Ma Jowina Galarion for general sequencing guidance, troubleshooting and help with setting up the sequencing laptop. We would also like to thank all members of the Streicker group for their feedback on the manuscript.

## Author Contributions

H.M. conceptualised the project; performed most of the formal analysis and investigation, including sample extractions, PCRs, library preparation, sequencing and data analysis; provided oversight of sample collection in Brazil; co-ordinated international sample shipments from other countries; and wrote the manuscript. E.F., A.C. & A.W. co-ordinated collection, storage and shipping of samples from Brazil. A.B. & M.G. inactivated, extracted and tested saliva swab samples from Peru. D.J.B., M.C.S., N.B.S, G.G.C. & S.S. co-ordinated collection, nucleic acid extraction and shipment of samples from Panamá and Belize. E.C. coordinated collection and shipment of samples from México. A.L. co-ordinated collection, nucleic acid extraction and shipping of samples from French Guiana. A.R. and C.C. tested Trinidad & Tobago *D. rotundus* samples, collected by N.M. and L.R., before co-ordinating shipment of amplicons to Glasgow for sequencing. R.O. wrote scripts for the bioinformatic processing of raw nanopore reads. C.T. coordinated field expeditions in Peru to obtain the samples used in this study. L.B. & M.J. provided project supervision and guidance. D.G.S. conceptualised, managed and supervised the project; acquired funding; and led review and editing of the manuscript, assisted by all other authors.

## Data accessibility statement

Datasets and phylogenetic trees used for analyses have been uploaded to FigShare (DOI: will receive once submitted). DrBHV DPOL-gB sequences will be uploaded to GenBank before publication.

## Code availability statement

R scripts used to generate data and figures and BEAST XML files have been uploaded to FigShare (DOI: will receive once submitted).

