## Supplementary Material for "Cross-Continental Analysis of Vampire Bat Betaherpesvirus Reveals Limited Interference Among Strains and Local Geographic Spread"

**For**

Haris Malik[1,2], Richard Orton[1], Anushka Ramjag[3], Fernando Gonçalves[4], Alice Broos[1], Aline Campos[5], Andre Witt[6], Anne Lavergne[7], Carlos Tello[8], Christine Carrington[3], Daniel Becker[9], Elsa Cardenas-Canales[10], Erich Fischer[11], Katherine Smollett[1], Mauro Galetti Rodrigues[12], Megan Griffiths[1], Sebastian Stockmaier[13], Ana Da Silva Filipe[1], Michael Jarvis[14], Laura Bergner[2], and Daniel G Streicker[1, 2]

[1]Medical Research Council–University of Glasgow Centre for Virus Research, Glasgow, United Kingdom
[2]School of Biodiversity, One Health and Veterinary Medicine, University of Glasgow, Glasgow, United Kingdom
[3]Department of Preclinical Sciences, Faculty of Medical Sciences, The University of the West Indies, St. Augustine, Republic of Trinidad and Tobago
[4]Department of Evolutionary Biology and Environmental Studies, University of Zurich, Zurich, Switzerland
[5]Centro Estadual de Vigilância em Saúde, Secretaria de Saúde do Estado do Rio Grande do Sul, Porto Alegre, RS, Brazil
[6]Secretaria Estadual de Agricultura, Pecuária e Desenvolvimento Rural (SEAPDR), Rio Grande do Sul, Porto Alegre 90150-004, Brazil
[7]Laboratoire des Interactions Virus-Hôtes, Institut Pasteur de la Guyane, Cayenne, French Guiana
[8]Association for the Conservation and Development of Natural Resources, Lima, Perú
[9]School of Biological Sciences, University of Oklahoma, Norman, Oklahoma, United States
[10]Department of Pathobiological Sciences, School of Veterinary Medicine, University of Wisconsin-Madison, Madison, Wisconsin, United States of America
[11]Instituto de Biociências, Universidade Federal de Mato Grosso do Sul, Campo Grande, MS 79070-900, Brazil
[12]Center for Biodiversity Dynamics and Climate Change (CBioClima), Department of Biodiversity, São Paulo State University (UNESP), Rio Claro, Sao Paulo, Brazil
[13]Department of Ecology and Evolutionary Biology, University of Tennessee, Knoxville, Tennessee, United Sates
[14] School of Biomedical Sciences, University of Plymouth; Devon, PL4 8AA, United Kingdom

**Contains:**

Supplementary Tables S1 → S6

Supplementary Figures S1 & S2

### Supplementary Tables

**Table S1: Top-ranked candidate binomial GLMMs explaining variation in DrBHV infection status.** Models were fitted in *glmmTMB* and ranked by corrected Akaike's Information Criterion (AICc) using *MuMIn::dredge*. The table shows the 10 most parsimonious candidate models. Candidate fixed effects included region, sex, age class, reproductive status, weight and forearm length, with capture year included as a random intercept. Because weight and forearm length were collinear body-size proxies, only one of these terms was permitted in any given model. Asterisks (*) indicate predictors for which at least one coefficient was significant at Pr(>|z|) < 0.05 within that model.

| Model | Sex | Weight | Repro Status | Age | Forearm | AICc | BIC |
|---|---|---|---|---|---|---|---|
| Amplification ~ Reproductive_Status + (1 \| Capture_Year) | | | * | | | 223.81 | 233.39 |
| Amplification ~ Forearm_z + Reproductive_Status + (1 \| Capture_Year) | | | * | | | 225.22 | 237.95 |
| Amplification ~ Reproductive_Status + Sex + (1 \| Capture_Year) | | | * | | | 225.69 | 238.42 |
| Amplification ~ Reproductive_Status + Weight_z + (1 \| Capture_Year) | | | * | | | 225.81 | 238.54 |
| Amplification ~ Age + Reproductive_Status + (1 \| Capture_Year) | | | * | | | 226.99 | 242.85 |
| Amplification ~ Forearm_z + Reproductive_Status + Sex + (1 \| Capture_Year) | | | * | | | 227.33 | 243.18 |
| Amplification ~ Reproductive_Status + Sex + Weight_z + (1 \| Capture_Year) | | | | | | 227.54 | 243.39 |
| Amplification ~ Age + Forearm_z + Reproductive_Status + (1 \| Capture_Year) | | | | | | 227.84 | 246.79 |
| Amplification ~ Age + Sex + (1 \| Capture_Year) | | | | | | 228.12 | 243.97 |
| Amplification ~ Sex + (1 \| Capture_Year) | | | | | | 228.14 | 237.72 |

**Table S2: Likelihood ratio test comparing models with and without the interaction between reproductive status and sex**. Outputs from the comparison of the most parsimonious GLMM (Reproductive Status) and another GLMM fitted with an interaction between Reproductive Status and Sex, using a Chi-square Likelihood Ratio Test (LRT). Capture year remained as a random effect.

| Model | AICc | BIC | Chisq | P value |
|---|---|---|---|---|
| Reproductive Status | 223.68 | 233.39 | NA | NA |
| Reproductive Status * Sex | 227.13 | 243.31 | 0.55 | 0.76 |

**Table S3: Additional Betaherpesvirus sequences from GenBank used to assess DrBHV monophyly.**

| Sequence ID | Host Species | GenBank Accession |
|---|---|---|
| FJ483970.2_AotineHerpesvirus1_strain_S34E | *Aotus trivirgatus* | FJ483970 |
| NC_020231.1_Caviid_herpesvirus_2_strain_21222 | *Cavia porcellus* | NC020231 |
| AY129397.2_ColobusGuereza_Cytomegalovirus | *Colobus guereza* | AY129397 |
| FJ538490_GorillaGorillaCMV | *Gorilla gorilla* | FJ538490 |
| KC465951.1_HHV-6A_GS | *Homo sapien* | KC465951 |
| KJ361971.1_HHV5_strain_UKNEQAS1 | *Homo sapien* | KJ361971 |
| KP745677.1_HHV5_strain_BE/1/2010 | *Homo sapien* | KP745677 |
| MT044477.1_HHV5_strain_GLA-SOT4 | *Homo sapien* | MT044477 |
| MW528463.1_HHV5_isolate_BM18 | *Homo sapien* | MW528463 |
| OK000909.1_HHV5_strain_Merlin | *Homo sapien* | OK000909 |
| AY28171_MacacaFascicularis_CMV | *Macaca fascicularis* | AY28171 |
| MT157323.1_CynomolgusCMV_strain31709 | *Macaca fascicularis* | MT157323 |
| DQ120516.1_MacacaMulatta_CMV | *Macaca mulatta* | DQ120516 |
| KX689268.1_Macacine_BHV_3_isolate_19936 | *Macaca mulatta* | KX689268 |
| NC_075417.1_MandrillusLeucophaeus_CMV_OCOM6-2 | *Mandrillus leucophaeus* | NC075417 |
| OP429138.1_MastomysNatalensis_CMV_1 | *Mastomys natalensis* | OP429138 |
| NC_076129.1_MiniopterusSchreibersii_HV1_B7D8 | *Miniopterus schreibersii* | NC_076129 |
| EU579859.2_Muromegalovirus_G4 | *Mus musculus* | EU579859 |
| EU579860.1_Muromegalovirus_WP15B | *Mus musculus* | EU579860 |
| EU579861.1_Muromegalovirus_C4A | *Mus musculus* | EU579861 |
| HE610454.1_Murid_herpesvirus_1_strain_N1 | *Mus musculus* | HE610454 |
| NC_003521.1_PanTroglodytesBHV_strain_Heberling | *Pan troglodytes* | NC003521 |
| FJ538485.2_PanTroglodytes_CMV | *Pan troglodytes* | FJ538485 |
| NC_003521.1_Panine_herpesvirus_2_strain_Heberling | *Pan troglodytes* | NC003521 |
| MT157322.1_Baboon_cytomegalovirus_strain_34826 | *Papio hamadryas* | MT157322 |
| NC027016_Papio_Ursinis_Cytomegalovirus_isolate_OCOM4-52 | *Papio ursinis* | NC027016 |
| AY129396_PongoPygmaeus_Cytomegalovirus | *Pongo pygmaeus* | AY129396 |
| OP429144.1_Murid_betaherpesvirus_2_strain_Maastricht | *Rattus norvegicus* | OP429144 |
| NC_002512.2_RatCMV_Maastricht | *Rattus rattus* | NC002512 |
| NC_002794.1_Tupaiid_herpesvirus_1 | *Tupaia spp* | NC002794 |

**Table S4: Well supported regional DrBHV transitions in Latin America.** Regional DrBHV transitions with BF support > 5. Some values have been rounded to 2 decimal places for viewing purposes. BF values were capped to 10000 to account for posterior inclusion probabilities of 1 (i.e. Inf posterior odds).

| From | To | Posterior Inclusion Prob | Prior Inclusion Prob | Posterior Odds | Prior Odds | Bayes Factor |
|---|---|---|---|---|---|---|
| Belize | Yucatan | 1.00 | 0.07 | Inf | 0.07143 | 10000 |
| Brazil | Cajamarca | 1.00 | 0.07 | Inf | 0.07143 | 10000 |
| Brazil | Cusco | 1.00 | 0.07 | Inf | 0.07143 | 10000 |
| Brazil | FrenchGuiana | 1.00 | 0.07 | Inf | 0.07143 | 10000 |
| Brazil | Huanuco | 1.00 | 0.07 | Inf | 0.07143 | 10000 |
| Brazil | Lima | 1.00 | 0.07 | Inf | 0.07143 | 10000 |
| Brazil | Trinidad | 1.00 | 0.07 | Inf | 0.07143 | 10000 |
| Cusco | Apurimac | 1.00 | 0.07 | Inf | 0.07143 | 10000 |
| Cusco | Ayacucho | 1.00 | 0.07 | Inf | 0.07143 | 10000 |
| Jalisco | Belize | 1.00 | 0.07 | Inf | 0.07143 | 10000 |

| | | | | | | |
|---|---|---|---|---|---|---|
| Jalisco | Yucatan | 1.00 | 0.07 | 844.57 | 0.07143 | 11824 |
| Brazil | Panama | 0.99 | 0.07 | 150.77 | 0.07143 | 2110.77 |
| Lima | Cajamarca | 0.99 | 0.07 | 102.84 | 0.07143 | 1439.79 |
| Brazil | Amazonas | 0.97 | 0.07 | 35.54 | 0.07143 | 497.52 |
| Brazil | Jalisco | 0.97 | 0.07 | 33.68 | 0.07143 | 471.54 |
| Panama | Belize | 0.96 | 0.07 | 24.12 | 0.07143 | 337.62 |
| Brazil | Loreto | 0.95 | 0.07 | 20.63 | 0.07143 | 288.8 |
| Panama | Lima | 0.95 | 0.07 | 20.45 | 0.07143 | 286.24 |
| Jalisco | Panama | 0.93 | 0.07 | 13.42 | 0.07143 | 187.95 |
| Yucatan | Belize | 0.89 | 0.07 | 8.02 | 0.07143 | 112.26 |
| Lima | Huanuco | 0.77 | 0.07 | 3.36 | 0.07143 | 46.98 |
| Huanuco | Amazonas | 0.61 | 0.07 | 1.57 | 0.07143 | 21.96 |
| Cajamarca | Huanuco | 0.38 | 0.07 | 0.60 | 0.07143 | 8.38 |
| Cajamarca | Amazonas | 0.37 | 0.07 | 0.59 | 0.07143 | 8.22 |
| Trinidad | Apurimac | 0.27 | 0.07 | 0.37 | 0.07143 | 5.23 |
| Trinidad | Ayacucho | 0.27 | 0.07 | 0.37 | 0.07143 | 5.16 |
| Cajamarca | Lima | 0.27 | 0.07 | 0.36 | 0.07143 | 5.08 |

**Table S5: BEAST GLM-CTMC model outputs.** Outputs are shown for both the external-transition and internal-transition models. BF values were capped to 10,000 to account for posterior inclusion probabilities of 1 (i.e. Inf posterior odds).

| Model | Predictor | Posterior Mean ± 95% HDI | Inclusion Prob | Prior Inclusion | Bayes Factor |
|---|---|---|---|---|---|
| External Transitions | Geodesic Distance | −0.995 [−1.268 to −0.708] | 1.000 | 0.5 | 10,000 |
| | Climate Similarity | 0.184 [−0.159 to 0.543] | 0.043 | 0.5 | 0.04 |
| | Sample Size (Origin) | 2.748 [1.403 to 4.135] | 1.000 | 0.5 | 10,000 |
| | Sample Size (Destination) | 0.080 [−0.290 to 0.371] | 0.027 | 0.5 | 0.03 |
| Internal Transitions | Geodesic Distance | −0.661 [−0.974 to −0.335] | 0.963 | 0.5 | 26.24 |
| | Climate Similarity | 0.242 [−0.103 to 0.546] | 0.151 | 0.5 | 0.18 |
| | Sample Size (Origin) | 3.873 [2.735 to 5.104] | 1.000 | 0.5 | 10,000 |
| | Sample Size (Destination) | 0.833 [0.436 to 1.237] | 0.997 | 0.5 | 319.57 |

**Table S6: Geographical coverage of DrBHV strains.**

| Strain | Region Count | Regions Found | Country Count | Countries Found |
|---|---|---|---|---|
| 1 | 1 | Brazil | 1 | Brazil |
| 2 | 4 | Amazonas, Cajamarca, Cusco, Trinidad | 2 | Peru, Trinidad |
| 3 | 4 | Apurimac, Cusco, FrenchGuiana, Trinidad | 3 | FrenchGuiana, Peru, Trinidad |
| 4 | 3 | Apurimac, Ayacucho, Trinidad | 2 | Peru, Trinidad |
| 5 | 4 | Belize, Jalisco, Lima, Yucatan | 3 | Belize, Mexico, Peru |
| 6 | 2 | Belize, Yucatan | 2 | Belize, Mexico |
| 7 | 3 | Amazonas, Cajamarca, Huanuco | 1 | Peru |

| | | | | |
|---|---|---|---|---|
| 8 | 1 | Brazil | 1 | Brazil |
| 9 | 1 | Brazil | 1 | Brazil |
| 10 | 1 | FrenchGuiana | 1 | FrenchGuiana |
| 11 | 1 | Trinidad | 1 | Trinidad |
| 12 | 2 | Cajamarca, Panama | 2 | Panama, Peru |
| 13 | 3 | Brazil, FrenchGuiana, Huanuco | 3 | Brazil, FrenchGuiana, Peru |
| 14 | 1 | Brazil | 1 | Brazil |
| 15 | 4 | Brazil, Cusco, FrenchGuiana, Loreto | 3 | Brazil, FrenchGuiana, Peru |
| 16 | 6 | Brazil, Cusco, FrenchGuiana, Lima, Loreto, Trinidad | 4 | Brazil, FrenchGuiana, Peru, Trinidad |
| 17 | 4 | Apurimac, Ayacucho, Brazil, Cusco | 2 | Brazil, Peru |
| 18 | 1 | Panama | 1 | Panama |
| 19 | 3 | Belize, Jalisco, Yucatan | 2 | Belize, Mexico |
| 20 | 2 | Belize, Jalisco | 2 | Belize, Mexico |
| 21 | 2 | Huanuco, Lima | 1 | Peru |
| 22 | 3 | Cajamarca, Huanuco, Lima | 1 | Peru |
| 23 | 1 | Jalisco | 1 | Mexico |
| 24 | 2 | Belize, Yucatan | 2 | Belize, Mexico |
| 25 | 1 | Jalisco | 1 | Mexico |
| 26 | 4 | Belize, Jalisco, Panama, Yucatan | 3 | Belize, Mexico, Panama |
| 27 | 1 | Jalisco | 1 | Mexico |
| 28 | 3 | Belize, Jalisco, Yucatan | 2 | Belize, Mexico |
| 29 | 2 | Jalisco, Yucatan | 1 | Mexico |
| 30 | 2 | Belize, Yucatan | 2 | Belize, Mexico |
| 31 | 1 | Jalisco | 1 | Mexico |
| 32 | 3 | Belize, Jalisco, Yucatan | 2 | Belize, Mexico |
| 33 | 1 | Belize | 1 | Belize |
| 34 | 1 | Jalisco | 1 | Mexico |
| 35 | 1 | Jalisco | 1 | Mexico |
| 36 | 2 | Lima, Panama | 2 | Panama, Peru |
| 37 | 1 | Panama | 1 | Panama |
| 38 | 3 | Belize, Panama, Yucatan | 3 | Belize, Mexico, Panama |
| 39 | 1 | Panama | 1 | Panama |
| 40 | 1 | Panama | 1 | Panama |
| 41 | 1 | Panama | 1 | Panama |
| 42 | 6 | Amazonas, Brazil, Cajamarca, Huanuco, Lima, Trinidad | 3 | Brazil, Peru, Trinidad |
| 43 | 3 | Brazil, Cusco, FrenchGuiana | 3 | Brazil, FrenchGuiana, Peru |
| 44 | 1 | Panama | 1 | Panama |
| 45 | 4 | Amazonas, Cajamarca, Lima, Trinidad | 2 | Peru, Trinidad |
| 46 | 2 | Brazil, FrenchGuiana | 2 | Brazil, FrenchGuiana |
| 47 | 2 | Brazil, FrenchGuiana | 2 | Brazil, FrenchGuiana |
| 48 | 3 | Amazonas, Brazil, Huanuco | 2 | Brazil, Peru |
| 49 | 4 | Amazonas, Brazil, FrenchGuiana, Huanuco | 3 | Brazil, FrenchGuiana, Peru |
| 50 | 2 | Brazil, FrenchGuiana | 2 | Brazil, FrenchGuiana |

##### Table S7: Model coefficients for strain co-circulation GLMM.

| Predictor | Estimate [95% CI: low → high] | p-value |
|---|---|---|
| Geodesic Distance to Nearest Region With Strain | -1.312 [95% CI: -1.718 → -0.906] | $2.42\times10^{-10}$ |
| Mean Genetic Distance to Regionally Circulating Strains | -0.445 [95% CI: -0.869 → -0.021] | 0.040 |
| Minimum Genetic Distance to Regionally Circulating Strains | -0.099 [95% CI: -0.556 → 0.357] | 0.670 |
| Sampling Effort (No. of Individuals/Region) | 0.328 [95% CI: -0.008 → 0.663] | 0.056 |
| Region Richness (No. of Strains/Region) | 0.669 [95% CI: 0.289 → 1.050] | 0.00056 |

#### Supplementary Figures

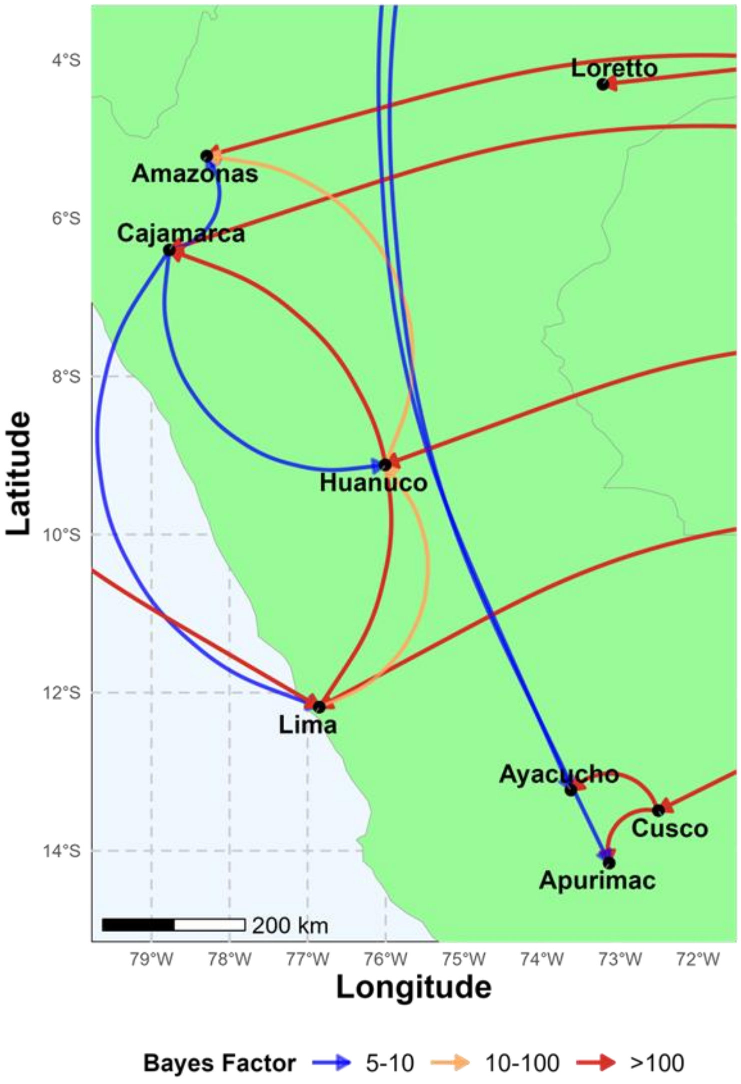


**Figure S1: Well-supported DrBHV Transitions in Peru.** Peruvian DrBHV transitions with BF support > 5. BF support was indicated by the colour of transition arrows. Red arrows from the right represent introductions from Brazil. The red arrow from the left represents an introduction from Panama. The two blue arrows represent two introductions from Trinidad and Tobago.

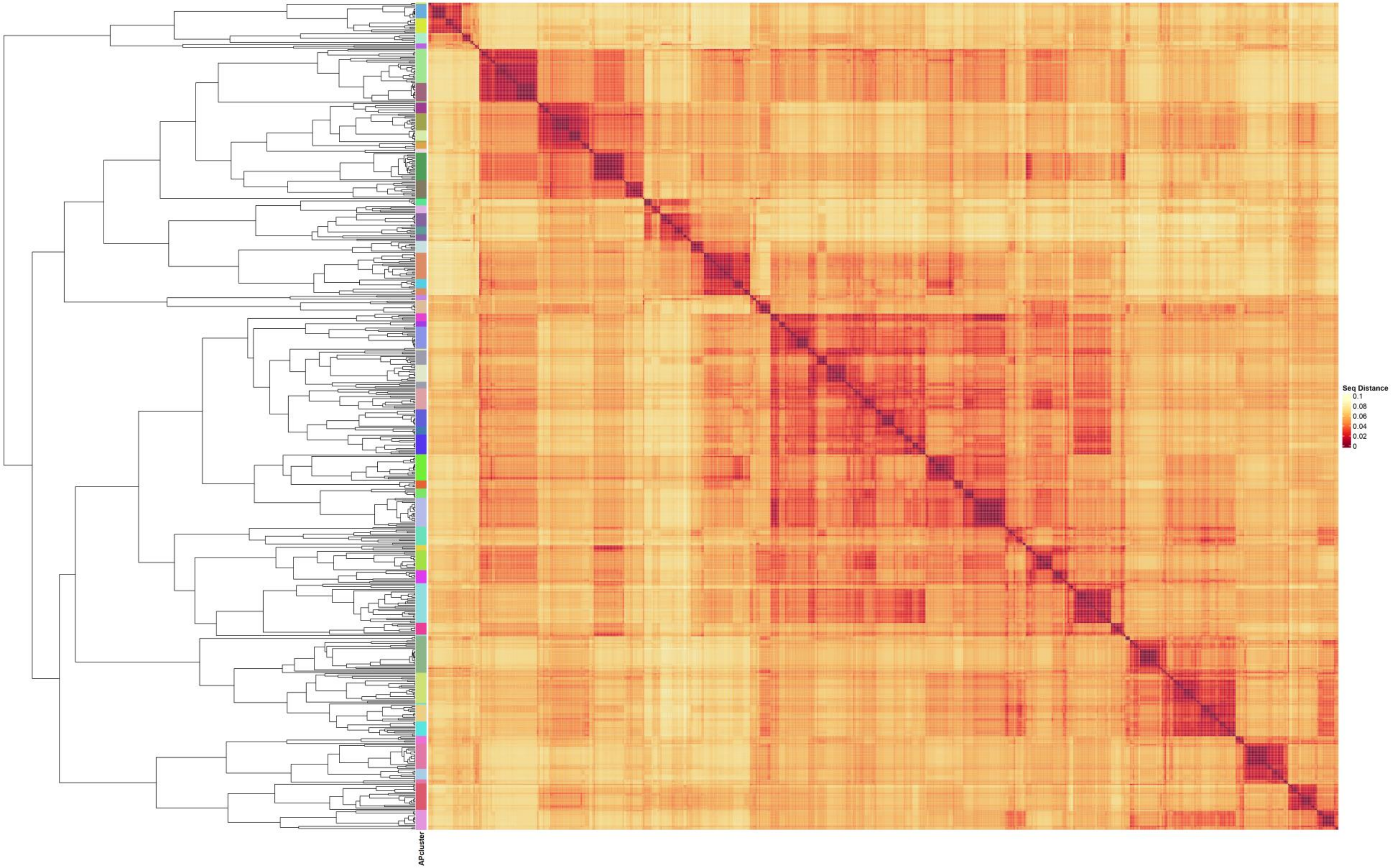


**Figure S2: Affinity propagation to delineate DrBHV strains.** Strains were delineated from the MCC tree using affinity propagation clustering (*apcluster* package in R, default settings). To assess the appropriateness of these strain boundaries, we also constructed a heatmap of pairwise nucleotide distances among sequences. Pairwise distances were calculated from the sequence alignment using *dist.dna()* with the "raw" model in the *ape* package.
